# Mesenchymal stem cell-derived extracellular vesicles attenuate symptoms in dextran sodium sulfate-induced ulcerative colitis mouse model

**DOI:** 10.1101/2024.02.01.578325

**Authors:** Yanhua Zhai, Guowu Liu, Chuang Cui, Yuan Yi, Yu Yan, Lei Zhang, Wan Wang, Xinjun He, Ke Xu

## Abstract

Inflammatory bowel disease (IBD) refers to a group of multifactorial and chronic inflammation affecting intestinal tract. Based on the pathogenic mechanisms, IBD mainly comprised of two categories: ulcerative colitis (UC) and Crohn’s disease (CD). In this study, we established a dextran sulfate sodium (DSS)-induced colitis model to mimic UC. Mesenchymal stem cell-derived extracellular vesicles (MSC-EVs) as therapeutic vectors have shown regenerating effect of damaged tissues. Here, intravenous administration of MSC-EVs ameliorated IBD symptoms including gaining weight, reducing disease activity index and restoring colon length. In addition, the protective effect of MSC-EVs were demonstrated by repairing colon mucosa and reducing the infiltration of macrophages in submucosa. Collectively, MSC-EV shows a therapeutic potential for IBD.

## Introduction

Inflammatory bowel disease (IBD), including ulcerative colitis and Crohn’s disease, is a chronic, relapsing gastrointestinal inflammatory disease of unknown etiology, with clinical symptoms including diarrhea, abdominal pain, fever, intestinal obstruction, etc. It is generally believed that IBD is caused by abnormal and exaggerated immune responses to environmental factors and gut microbes in genetically susceptible individuals, but its exact etiology remains to be further investigated^1^.

Cytokine hypersecretion and chronic inflammation are typical features of IBD, which are caused by an imbalance between inflammatory, regulatory, and anti-inflammatory cytokines. The expression of pro-inflammatory factors (such as TNF-α, IL-6, and IL-23) is significantly increased in IBD patients. These cytokines directly or indirectly cause intestinal epithelial cell damage or necrosis, thus promoting the occurrence and development of IBD. Inhibitors targeting TNF-α (infliximab, adalimumab, golimumab, and certolizumab pegol), IL-12 and IL-23 (Ustekinumab), and JAK (Tofacitinib) have been approved by FDA for clinical treatment of IBD^2^. Both innate immune cells and adaptive immune cells are involved in the abnormal immune response of IBD^3^. Innate immune cells such as macrophages, dendritic cells, neutrophils and NKT are involved in the disease process of IBD. Macrophages and dendritic cells express pattern recognition receptors (PRRs), such as Toll-like receptors (TLRs) and NOD-like Receptors (NLRs). When pathogen-associated molecular patterns (PAMPs) bind to these receptors, a variety of intracellular signaling pathways are activated to promote the secretion of inflammatory factors and chemokines. In addition, macrophages and dendritic cells also present antigens to T cells to activate adaptive immunity. Macrophages are considered potential new targets for developing new treatments for IBD^4,5^.

Adaptive immune cells have high specificity and immune memory capabilities, and the key players are T cells. Research has found that Th1/Th17 is closely related to Crohn’s disease. Th1 cells secrete IFN-γ, TNF-α and IL-2, and recruit macrophages, NK cells and CD8+ T cells. TNF-α can cooperate with IFN-γ to kill intestinal epithelial cells and destroy the intestinal epithelial barrier function^6^. Th17 cells produces cytokines, such as IL-17A, IL-17F, IL-21, and IL-22, which are important drivers of inflammation in IBD. Clinical studies have found that the levels of Th17 cells, IL-17, and IL-23 are significantly elevated in the intestines of IBD patients ^7^.

Mesenchymal stem cells (MSCs) exist in various tissues such as bone marrow, umbilical cord tissue and adipose tissue, and have multi-directional differentiation potential. A large number of studies have shown that MSCs have great potential in the treatment of various diseases, including osteoarthritis, pulmonary fibrosis, autoimmune diseases, organ transplantation, etc., by regulating inflammatory responses and participating in tissue repair and regeneration. When stimulated by inflammatory factors, MSCs will produce a large number of immune regulatory factors, chemokines and growth factors, thereby regulating the tissue immune microenvironment and promoting tissue regeneration. Researchers have found that MSCs exert therapeutic effects mainly by secreting cytokines (growth factors and chemokines) and extracellular vesicles (EVs). EVs are essential paracrine effectors of mesenchymal stem cells. Compared with mesenchymal stem cells, MSC-EVs have broader therapeutic application prospects and advantages: excellent biocompatibility and lower immunogenicity, ability for drug loading, better safety and easy storage for a long time. Therefore, using MSC-EVs to replace mesenchymal stem cells will be the focus of future clinical treatments. A large number of studies have shown that in DSS-induced mouse IBD models, extracellular vesicles secreted by bone marrow mesenchymal stem cells (BMSCs) are taken up by macrophages, thereby inhibiting the NF-κB signaling pathway and promoting M2 macrophage polarization, reducing the expression levels of inflammatory factors such as TNF-α and IL-12, and increasing the levels of IL-10 and TGF-β to relieve intestinal inflammation^8-10^. In addition, MSC-EVs derived from adipose tissue and human umbilical cord also have significant therapeutic effects on IBD^11,12^.

## Results

### MSC-EVs ameliorated the severity of colitis symptoms in the established DSS-induced mouse model

Therapeutic effects of MSC-EVs on IBD were explored through DSS-induced colitis. We established an optimized IBD mouse model by administrating DSS at a concentration of 2.5% (w/v) in drinking water for 8 days, MSC-EVs (200 μg) were administrated intravenously at day 3, 5 and 7 **(Figure 1)**.

**Figure 1.**
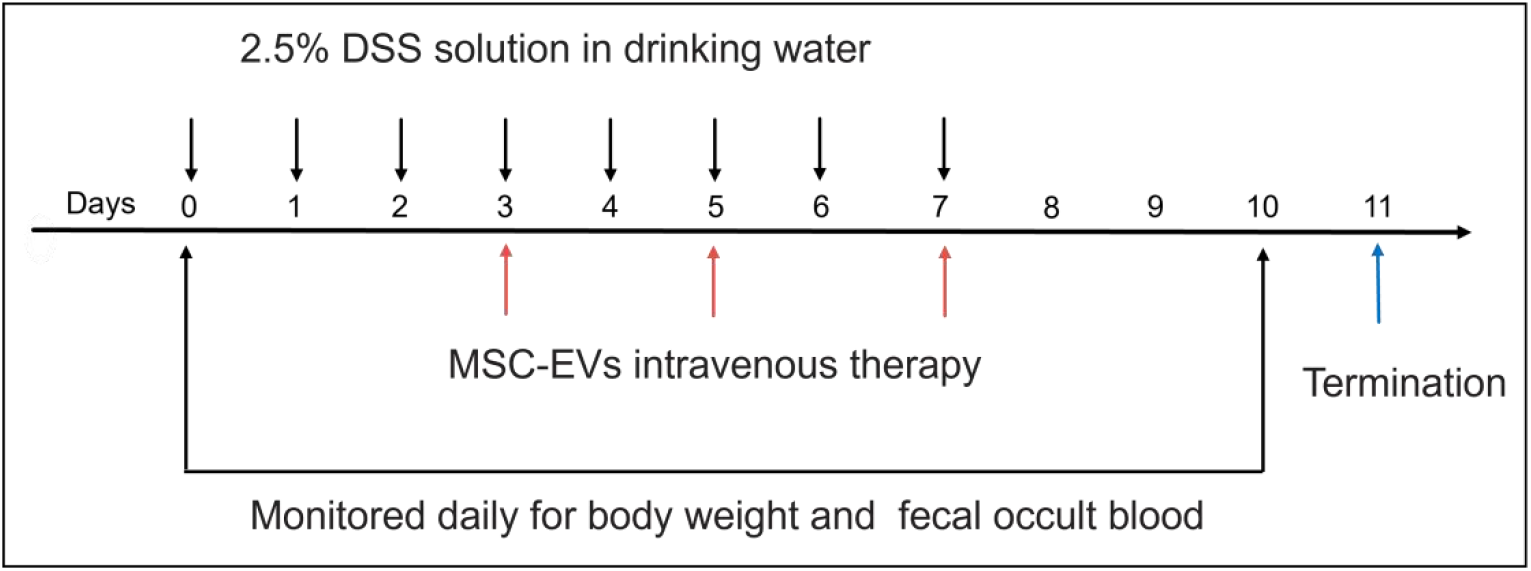
Schematic illustration of the experimental design. Mice were weighed and randomized into three groups: Un-Treated group with regular drinking water; DSS group with oral administration of 2.5% DSS, MSC-EVs group with intravenous administration of MSC-EVs for ameliorating symptoms of 2.5% DSS inducing colitis. 2.5% (w/v) DSS was supplied for 8 consecutive days from day 0 to day 7, MSC-EVs of 200 μg total protein were administrated intravenously per mouse every other day at day3, 5, 7. All mice in three groups were monitored for body weight and collected feces for occult blood and stool consistency analyses daily. On the day 11 of euthanasia, spleens, colons and ceca were isolated for analysis.

DSS lowered body weights **(Figure 2A)** and increased DAI scores **(Figure 2D)** compared to untreated mice. Meanwhile, fecal occult blood and stool were measured daily via comparative analysis. DSS treated mice showed soft or watery stools, accompanied by occult blood, while MSC-EVs administration significantly alleviated these symptoms **(Figure 2B and 2C)**. In line with these results, MSC-EVs ameliorated colon length reduction in DSS-induced mice on the day of termination **(Figure 2E and 2F)**.

**Figure 2.**
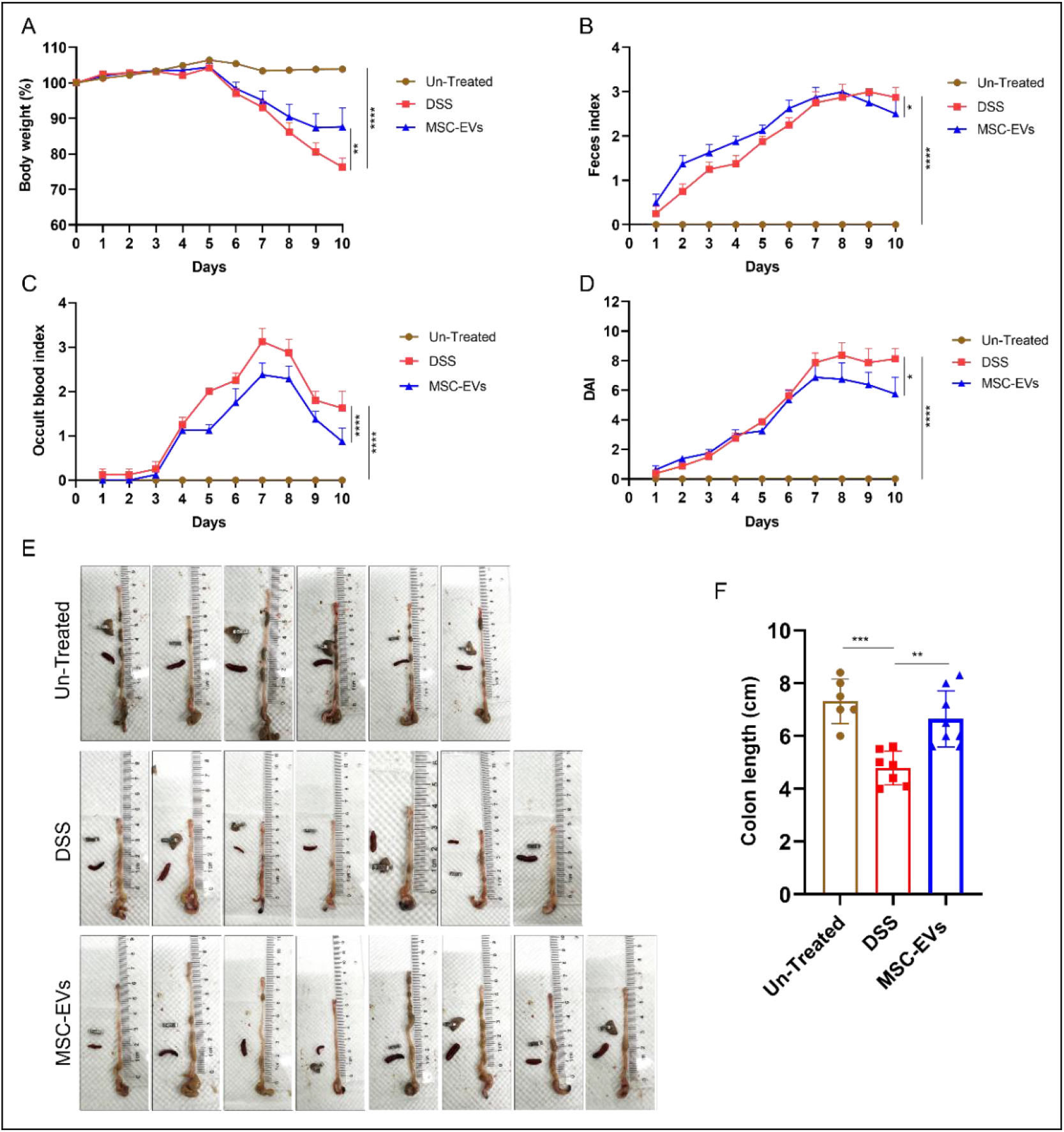
MSC-EVs alleviated DSS-induced colitis. (A) Body weight loss of mice with different treatment was recorded daily from day 0 to day 10. Feces was collected daily from day 0 to day 10 to monitor (B) stool consistency and (C) occult blood. (D) Disease activity index (DAI) score was decided by body weight change, occult blood and stool consistency. (E) Colon length of mice with different treatment was measured on the day of termination, with spleen and ear tag for comparison. (F) The statistical analysis of colon length was shown in different groups. Data were presented as mean±SEM. \**P*<0.05, \*\**P*<0.01, \*\*\**P*<0.001, \*\*\*\**P*<0.0001.

### MSC-EVs regenerated histological injury in DSS-induced colitis

Colon tissues from different groups of mice were analyzed for further histology. The sagittal sections of untreated mice showed a normal structure, whereas severe colonic mucosa damage was observed in DSS-induced mice, as characterized by damaged mucosal structure, such as distorted crypt and villus architecture, depleted goblet cells and intestinal glands, mucosal ulceration and erosion. Correspondingly, DSS induced an influx of immune cells into mucosa and submucosa. Further, histological analysis of colonic mucosa detected ulcer formation was decreased and colon morphology was massively improved through MSC-EVs treatment **(Figure 3)**. Furthermore, MSC-EVs administration notably alleviated inflammatory infiltration further revealed that the MSC-EVs treatment in IBD relative to the immunological regulation.

**Figure 3.**
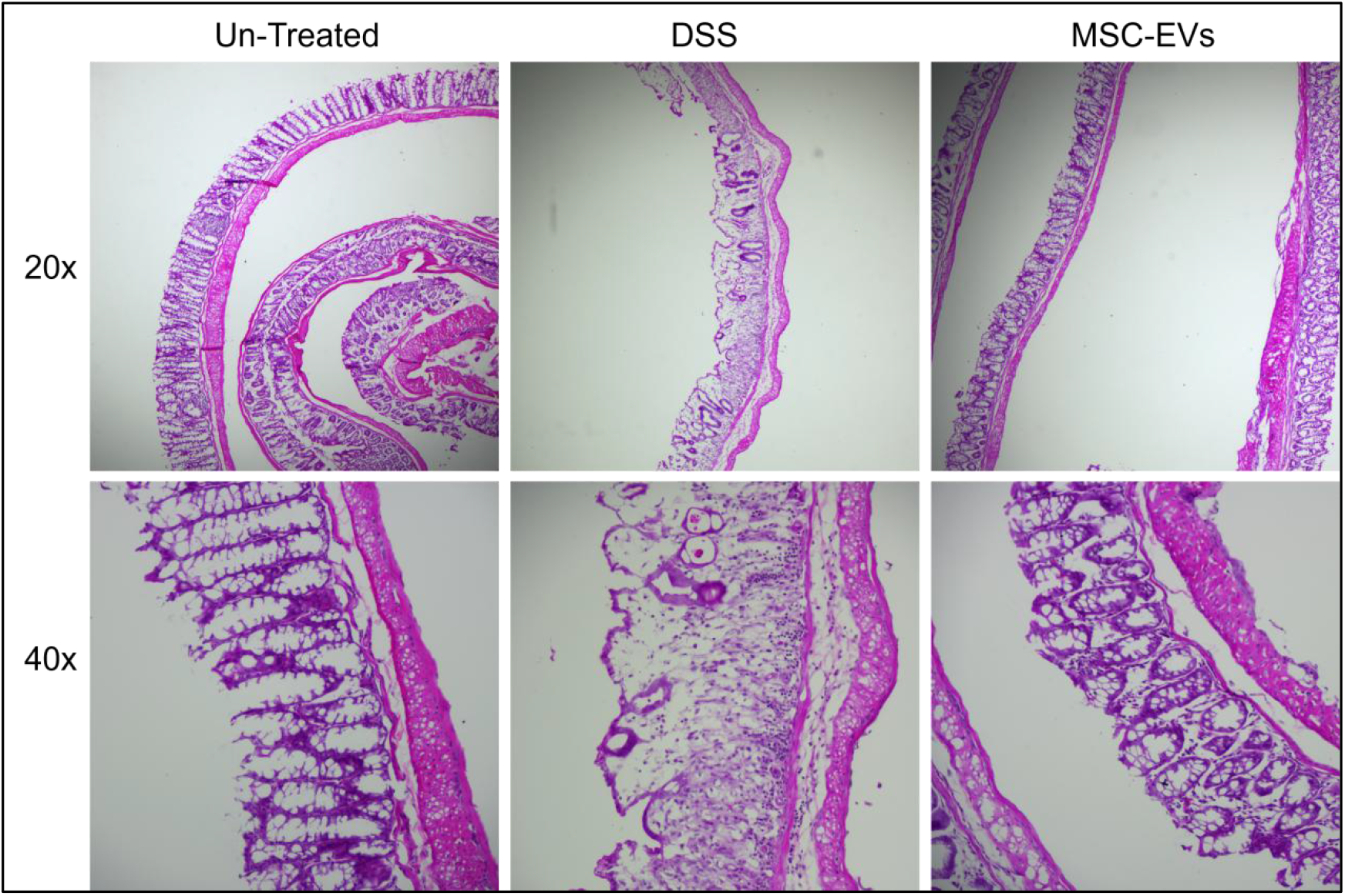
H&E-stained colonic sections showed MSC-EVs efficiently ameliorated morphological integrity of colons treatment with DSS. Untreated mice showed normal structures, DSS-induced mice had severe mucosal damage, whereas MSC-EVs significantly restored mucosal damage in mice with DSS-induced colitis. Images were taken at 20× and 40× magnification.

### MSC-EVs regulated the redistribution of macrophages in DSS-induced colitis

Immune dysfunction remains important pathogenesis of IBD. We future explored the inflammatory response of innate immune cells such as macrophages. DSS-treated mice showed a decreased expression of macrophage marker F4/80 in colonic mucosa and an increased expression in submucosa, whereas MSC-EVs administration effectively decreased the macrophage infiltration in submucosa. Additionally, MSC-EVs maintained macrophage level in DSS-induced colitis compared to untreated mice **(Figure 4)**. These observations all suggested that MSC-EVs exerted anti-inflammatory effects by restored the macrophage infiltration in colonic mucosa.

**Figure 4.**
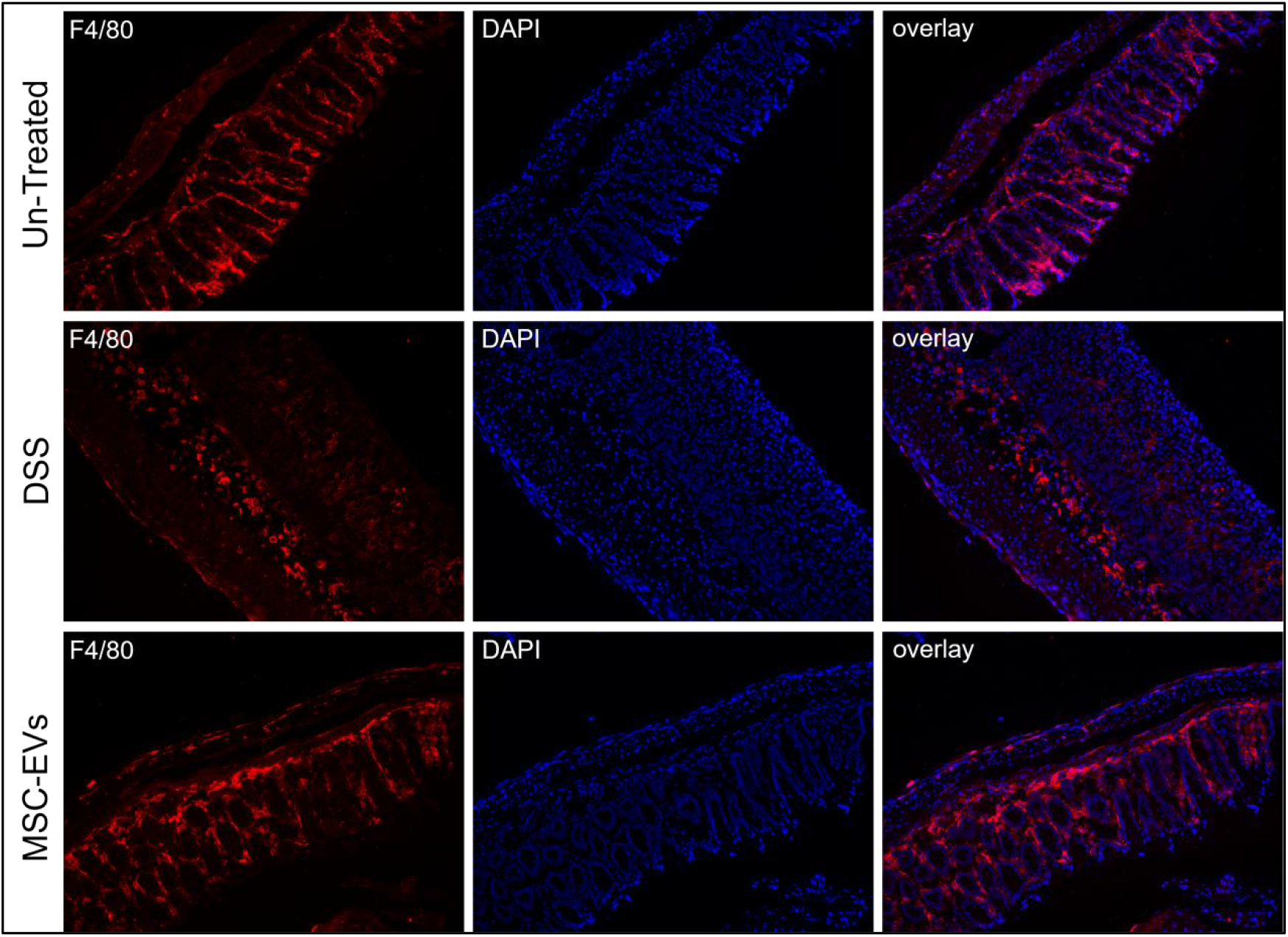
MSC-EVs alleviated macrophage infiltration in submucosa. Immunofluorescence labeling of macrophage marker F4/80 showed macrophages were mainly distributed in mucosa in untreated mice and MSC-EVs treated mice with DSS-induced colitis, whereas macrophages in mice with DSS treatment were widely distributed in submucosa. Images were taken at 20× magnification.

## Discussion

MSC-derived EVs have exerted therapeutic effects by repairing and regenerating tissue injuries. In this study, we aimed to test if MSC-EVs ameliorate DSS-induced colitis. According to our experimental data, DAI score, body weight, occult blood and stool consistency proved that MSC-EVs alleviated symptoms of DSS-induced colitis in mice. We observed typical histological changes induced by DSS include mucosa layer damage, whereas administration of MSC-EVs exerted a protective effect on mucosa. In addition, we explored that DSS could affect colonic immune response through macrophages infiltration in submucosa, MSC-EVs reduced the influx of macrophages into submucosa layer. Our study expanded the understanding of MSC-EVs in DSS-induced colitis treatment and provided a significant step for the preclinical research on IBD therapy.

## Materials and Methods

### Isolation and identification of extracellular vesicles

Upon culture termination, the cell culture supernatant was first centrifuged at 120 g at 4 °C for 5 min to remove cell precipitation, then clarified supernatant was transferred and centrifuged for 30 min at 16,000 g at 4 °C to remove large cell debris. The supernatant was transferred again and centrifuged at 133,900 g at 4 °C at 60 min to isolate EVs, the EVs were finally resuspended in PBS.

EV solution was diluted with PBS and measured total protein using pierce micro BCA protein assay kit (ThermoFisher) according to the manufacturer’s protocol. EV solution was diluted in PBS to a final volume of 1 mL, size distributions and concentrations were measured by nanoparticle tracking analysis (NTA) on a ZetaView (ParticleMetrix), all settings were set according to the manufacturer’s software manual.

### DSS induced ulcerative colitis mouse model

Prepared 2.5% (w/v) DSS (MP Biomedicals) solution with autoclaved drinking water, filled the cage water bottle and changed with freshly prepared DSS solution every 2 days. Female, 8-weeks-old C57BL/6J mice (Vital River) were administrated 2.5% DSS by oral intake, control mice got the same drinking water without DSS. DSS solution was supplied for 8 consecutive days (D0-D7). Intravenous injection of MSC-EV was performed on day 3, 5 and 7. Body weight and occult blood were measured daily from D0 to D10. On day11, mice were euthanized in accordance with approved institutional animal ethical protocols, colons were isolated from ceca and the length of colons were measured. In addition, disease activity index was calculated through body weight change, occult blood and stool consistency.

### Histological staining

Colons were separated from the ceca and cut longitudinally, fixed as swiss rolls in 4% PFA, dehydrated in 30% sucrose solution, embedded with cryoprotective medium OCT and sectioned. Histopathological staining was performed with Hematoxylin & Eosin (H&E) staining kit (Beyotime) according to manufacturer’s protocol. For immunofluorescence staining, primary antibodies against F4/80 (Abcam) were incubated overnight at 4°C and the secondary antibodies (Invitrogen) were incubated for 1 h at room temperature.

### Statistical analysis

All experimental data were expressed as mean ± SEM and analyzed using GraphPad Prism 8 software. One-way analysis of variance (ANOVA) followed by Tukey’s multiple comparisons test was performed to compare colon length data. All other data were compared by Two-way ANOVA with Dunnett’s multiple comparisons test. P<0.05 was considered statistically significant.

